# An intranasal vaccine durably protects against SARS-CoV-2 variants in mice

**DOI:** 10.1101/2021.05.08.443267

**Authors:** Ahmed O. Hassan, Swathi Shrihari, Matthew J. Gorman, Baoling Ying, Dansu Yuan, Saravanan Raju, Rita E. Chen, Igor P. Dmitriev, Elena Kashentseva, Lucas J. Adams, Pei-Yong Shi, Daved H. Fremont, David T. Curiel, Galit Alter, Michael S. Diamond

## Abstract

SARS-CoV-2 variants that attenuate antibody neutralization could jeopardize vaccine efficacy and the end of the COVID-19 pandemic. We recently reported the protective activity of a single-dose intranasally-administered spike protein-based chimpanzee adenovirus-vectored vaccine (ChAd-SARS-CoV-2-S) in animals, which has advanced to human trials. Here, we assessed its durability, dose-response, and cross-protective activity in mice. A single intranasal dose of ChAd-SARS-CoV-2-S induced durably high neutralizing and Fc effector antibody responses in serum and S-specific IgG and IgA secreting long-lived plasma cells in the bone marrow. Protection against a historical SARS-CoV-2 strain was observed across a 100-fold vaccine dose range and over a 200-day period. At 6 weeks or 9 months after vaccination, serum antibodies neutralized SARS-CoV-2 strains with B.1.351 and B.1.1.28 spike proteins and conferred almost complete protection in the upper and lower respiratory tracts after challenge. Thus, in mice, intranasal immunization with ChAd-SARS-CoV-2-S provides durable protection against historical and emerging SARS-CoV-2 strains.

## INTRODUCTION

Severe acute respiratory syndrome coronavirus 2 (SARS-CoV-2) is the etiologic agent of the Coronavirus Disease 2019 (COVID-19) syndrome, which can rapidly progress to pneumonia, respiratory failure, and systemic inflammatory disease ^1, 2, 3^. The elderly, immunocompromised, and those with certain co-morbidities (*e*.*g*., obesity, diabetes, and hypertension) are at greatest risk of severe disease, requirement of mechanical ventilation, and death ^4^. More than 152 million infections and 3.2 million deaths have been recorded worldwide (https://covid19.who.int) since the start of the pandemic. The extensive morbidity and mortality associated with COVID-19 pandemic have made the development and deployment of SARS-CoV-2 vaccines an urgent global health priority.

The spike (S) protein of the SARS-CoV-2 virion is the principal target for antibody-based and vaccine countermeasures. The S protein serves as the primary viral attachment and entry factor and engages the cell-surface receptor angiotensin-converting enzyme 2 (ACE2) to promote SARS-CoV-2 entry into human cells ^5^. SARS-CoV-2 S proteins are cleaved to yield S1 and S2 fragments ^6^, with the S1 protein containing the receptor binding domain (RBD) and the S2 protein promoting membrane fusion and virus penetration into the cytoplasm. The prefusion form of the SARS-CoV-2 S protein ^7^ is recognized by potently neutralizing monoclonal antibodies ^8, 9, 10, 11, 12^ or protein inhibitors ^13^.

Many vaccine candidates targeting the SARS-CoV-2 S protein have been developed ^14^ using DNA plasmid, lipid nanoparticle encapsulated mRNA, inactivated virion, protein subunit, or viral-vectored vaccine platforms ^15^. Several vaccines administered by intramuscular (IM) injection (*e*.*g*., Pfizer/BioNTech BNT162b2 and Moderna 1273 mRNA ^16, 17^ and Johnson & Johnson Ad26.COV2 and AstraZeneca ChAdOx1 nCoV-19 adenoviral ^18, 19^ platforms) have been granted emergency use authorization in many countries with hundreds of million of doses given worldwide (https://covid19.who.int).

While vaccines administered by IM injection induce robust systemic immunity that protects against severe disease and mortality, questions remain as to their ability to curtail SARS-CoV-2 transmission, especially if upper airway infection is not reduced. Indeed, many of the IM-administered vaccines showed variable protection against upper airway infection and transmission in pre-clinical studies and failed to induce substantive mucosal (IgA) immunity ^20, 21, 22, 23^. This issue is important because of the emergence of more transmissible SARS-CoV-2 variants including B.1.1.7, B.1.351, and B.1.1.28 with substitutions in the spike protein. Experiments with pseudoviruses and authentic SARS-CoV-2 strains also suggest that neutralization by vaccine-induced sera is diminished against variants expressing mutations in the spike gene at positions L452, E484, and elsewhere ^24, 25, 26, 27, 28, 29^. Beyond possible negative impacts on protection, the combination of diminished immunity against certain variants and naturally lower anti-S IgG levels in the respiratory mucosa could create conditions for further selection of resistance in the upper airway and transmission into the general population.

We recently described a single-dose, intranasally (IN)-delivered chimpanzee Adenovirus (simian Ad-36)-based SARS-CoV-2 vaccine (ChAd-SARS-CoV-2-S) encoding a pre-fusion stabilized S protein that induced robust humoral, cell-mediated, and mucosal immune responses and limited upper and lower airway infection in K18-hACE2 transgenic mice, hamsters, and non-human primates ^30, 31, 32^. This vaccine, which has advanced to human clinical trials (BBV154, Clinical Trial NCT04751682), differs from ChAdOx1 nCoV-19, a chimpanzee Ad-23-based SARS-CoV-2 vaccine, currently granted emergency use in some countries. Here, as a further step to evaluating the potential utility of ChAd-SARS-CoV-2-S, we assessed its dose-response, durability, and cross-protective activity in mice including effects on upper and lower airway infection. At approximately nine months after IN immunization, neutralizing antibody and anti-S protein IgA levels in serum of ChAd-SARS-CoV-2-S-vaccinated animals remained high and inhibited infection with SARS-CoV-2 strains with B.1.351 and B.1.1.28 spike proteins. At this time, susceptible K18-hACE2 transgenic mice were fully protected against upper and lower respiratory tract infection after challenge with a SARS-CoV-2 virus displaying B.1.351 spike proteins.

## RESULTS

### A single ChAd-SARS-CoV-2-S immunization induces durable anti-spike and neutralizing responses at different doses

We assessed the durability of humoral immune responses in BALB/c mice 100 or 200 days post IM-or IN-immunization with escalating doses of ChAd-SARS-CoV-2-S (10^8^, 10^9^, and 10^10^ viral particles [vp]) or 10^10^ vp of a ChAd-Control vaccine (**Fig 1A**). First, we measured anti-S and anti-RBD IgG and IgA levels by ELISA. Consistent with prior results at a one-month time point ^30^, at 100 or 200 days post-vaccination, IN immunization with ChAd-SARS-CoV-2-S induced superior antibody responses than IM immunization or vaccination with ChAd-Control (**Fig 1B-M and Fig S1**). Anti-S and anti-RBD-specific binding IgG levels in serum were greater after IN than IM immunization at 100 or 200 days. At 100 days after IN immunization with 10^10^, 10^9^, and 10^8^ vp of ChAd-SARS-CoV-2-S, geometric mean titers (GMT) of S-specific IgG responses were 1.1 × 10^6^, 4.8 × 10^5^, and 2.6 × 10^5^, and RBD-specific IgG were 3.2 × 10^5^, 1.8 × 10^5^, and 8.7 × 10^4^, respectively (**Fig 1B**). In comparison, S- and RBD-specific IgG reponses 100 days after IM immunization with 10^10^, 10^9^, and 10^8^ vp of ChAd-SARS-CoV-2-S were 4 to 6-fold lower (*P* < 0.0001) with S-specific IgG titers of 2.1 × 10^5^, 1.1 × 10^5^, and 4.5 × 10^4^ and RBD-specific IgG titers of 5.1 × 10^4^, 2.9 × 10^4^, and 2.3 × 10^4^, respectively (**Fig 1E**). A similar dose response was observed with S-and RBD-specific IgG titers at 200 days after IN or IM immunization (**Fig 1H and K**). At 200 days after IN immunization with 10^10^, 10^9^, and 10^8^ vp of ChAd-SARS-CoV-2-S, GMT of S-specific IgG were 2.8 × 10^6^, 2.4 × 10^6^, and 1.2 × 10^6^, and RBD-specific IgG were 1.1 × 10^6^, 6.1 × 10^5^, and 3.2 × 10^5^, respectively (**Fig 1H**). At 200 days after IM immunization with 10^10^, 10^9^, and 10^8^ vp of ChAd-SARS-CoV-2-S, S-specific IgG GMT were 8.1 × 10^5^, 6.9 × 10^5^, and 2.6 × 10^5^, and RBD-specific IgG GMT were 1.4 × 10^5^, 1.3 × 10^5^, and 8.0 × 10^4^, respectively (**Fig 1K**). Thus, anti-S and anti-RBD IgG levels were higher after IN than IM immunization and continued to rise in serum even several months after single-dose vaccination.

**Figure 1.**
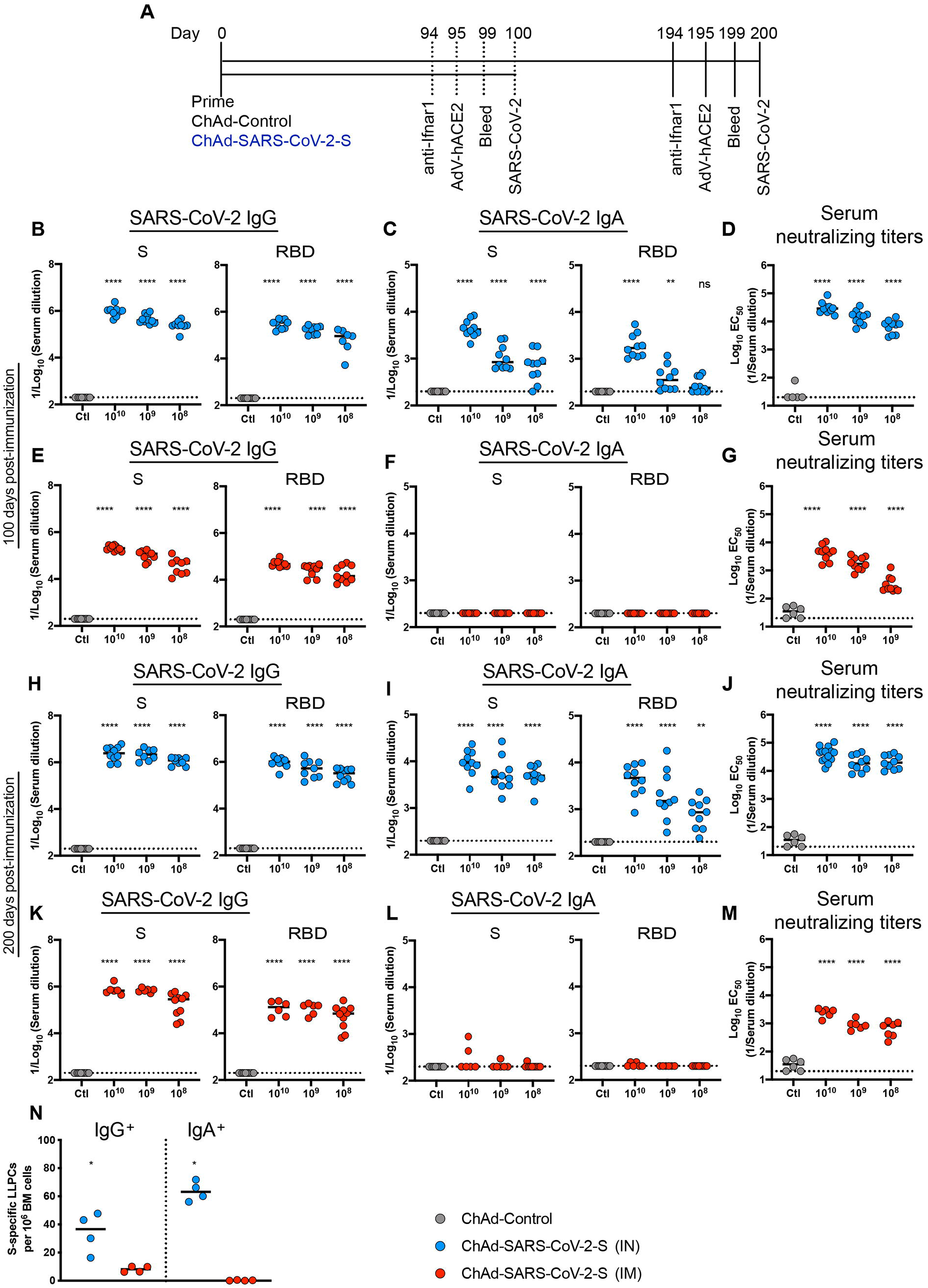
ChAd-SARS-CoV-2-S induces durable immunity. **A**. Immunization scheme. Five-week old female BALB/c mice were vaccinated via IN or IM route with 10^10^ viral particles of ChAd-control or decreasing doses (10^10^, 10^9^ and 10^8^ vp) of ChAd-SARS-CoV-2-S. **B-M**. Humoral responses in sera of immunized mice were evaluated (n = 6-14). An ELISA measured anti-S and RBD IgG and IgA levels from IN-immunized mice at 100 (**B-C**) or 200 (**H-I**) days post-vaccination, or from IM-immunized mice at 100 (**E-F**) or 200 (**K-L**) days post-vaccination. Neutralizing activity of sera determined by FRNT from IN-(**D, J**) or IM-(**G, M**) immunized mice at 100 (**D, G**) or 200 (**J, M**) days post-vaccination. **N**. The frequency of S-specific IgG or IgA producing LLPCs in the bone marrow measured by ELISPOT assay (n = 4). For (**B-M**): one way ANOVA with a Dunnett’s post-test comparing vaccine and control groups: ns, not significant; **, *P* < 0.01; ****, *P* < 0.0001). For (**N**): Mann-Whitney test: *, *P* < 0.05. **B-N**, Bars show median values, and dotted lines indicate the limit of detection (LOD) of the assays.

We next assessed the induction and durability of serum IgA responses. Although IM immunization failed to induce S- or RBD-specific IgA (**Fig 1F and L**), substantial levels of anti S- and RBD IgA were detected after IN immunization at 100 or 200 days post-immunization (**Fig 1C and I**). At 100 days after IN immunization with 10^10^, 10^9^, and 10^8^ vp of ChAd-SARS-CoV-2-S, the GMT of S-specific IgA were 4.8 × 10^3^, 1.2 × 10^3^, and 8.4 × 10^2^, and RBD-specific IgA were 2.2 × 10^3^, 4.6 × 10^2^, and 2.9 × 10^2^, respectively (**Fig 1C**). As seen with IgG, the IgA levels continued to increase over time such that at 200 days the GMT of S-specific IgA were 1.1 × 10^4^, 7.4 × 10^3^, and 5.4 × 10^3^, and RBD-specific IgA were 5.2 × 10^3^, 3.8 × 10^3^, and 9.8 × 10^2^ after IN immunization with 10^10^, 10^9^, and 10^8^ vp of ChAd-SARS-CoV-2-S, respectively (**Fig. 1I**).

We next evaluated a functional correlate of the serological response by assaying neutralizing activity (**Fig 1D, G, J, M and S1**) using a focus-reduction neutralization test (FRNT) ^33^. As expected, neutralizing activity was not detected in sera from ChAd-control treated mice. At 100 days post IN immunization with 10^10^, 10^9^, and 10^8^ vp of ChAd-SARS-CoV-2-S, the mean effective half maximal inhibitory titers [EC50] were 39,449, 9,989, and 7,270, respectively (**Fig 1D**). In comparison, at this time point after IM immunization with 10^10^, 10^9^, and 10^8^ vp of ChAd-SARS-CoV-2-S, the EC50 values were 8 to 20-fold lower (*P* < 0.0001) at 4,988, 2,017, and 391, respectively (**Fig 1G**). At 200 days after IN immunization with 10^10^, 10^9^, and 10^8^ vp of ChAd-SARS-CoV-2-S, and consistent with the higher anti-S and RBD titers seen, EC50 values were 45,591, 22,769, and 23,433, respectively (**Fig 1J**). In comparison, 200 days after IM immunization with 10^10^, 10^9^, and 10^8^ vp of ChAd-SARS-CoV-2-S, EC50 values were much lower at 2,524, 940, and 716, respectively (**Fig 1M**).

Long-lived plasma cells (LLPCs) reside in the bone marrow and constitutively secrete high levels of antibody that correlate with serum levels ^34^. To assess the levels of antigen-specific LLPCs at 200 days after IM or IN immunization with 10^10^ vp of ChAd-SARS-CoV-2-S, CD138^+^ cells were isolated from the bone marrow and assyed for S-specific IgG or IgA production using an ELISPOT assay ^35^. We observed a ∼4-fold higher frequency of LLPCs secreting S-specific IgG after IN immunization than IM immunization (**Fig 1N**). Additionally, after IN immunization, we detected higher numbers of LLPCs producing S-specific IgA, which were absent after IM immunization (**Fig 1N**). Together, these data establish the following: (a) single-dose IN immunization promotes superior humoral immunity than IM immunization; (b) 100-fold lower inoculating doses of ChAd-SARS-CoV-2-S induce robust neutralizing antibody responses in mice; (c) IN but not IM immunization induces serum IgA responses and IgA-specific LLPCs against the SARS-CoV-2 S protein; and (d) the humoral immunity induced by ChAd-SARS-CoV-2-S is durable and rises over a six-month period after vaccination.

### IN inoculation of ChAd-SARS-CoV-2-S induces broad antibody responses with Fc effector function capacity

To characterize the humoral response further, we analyzed antibody binding to SARS-CoV-2 variant proteins and Fc effector functions using serum derived from BALB/c mice at 90 days after IN or IM vaccination. Our panel of SARS-CoV-2 proteins included spike (D614G, E484K, N501Y, Δ69-70, K417N) and RBD (E484K) antigens corresponding to WA1/2020, B.1.1.7, B.1.351, B.1.1.28 strains. We first measured the anti-SARS-CoV-2 specific antibody response for several isotypes (IgG1, IgG2a, IgG2b, IgG3, IgM, and IgA) and their ability to bind Fcγ receptors (mouse FcγRIIB, FcγRIII, FcγRIV) using a luminex platform. Consistent with data obtained by ELISA (**Fig 1B and E**), IN vaccination of ChAd-SARS-CoV-2-S induced higher levels of IgG1 to D614G spike and WA1/2020 RBD proteins than IM immunization, and as expected, decreasing doses of the vaccine elicited lower antibody titers (**Fig 2A**). Anti-SARS-CoV-2 IgG1 titers after IN immunization also were higher against all spike and RBD variants than after IM immunization, and titers decreased with vaccine dose (**Fig 2B**). As shown in a heatmap, this trend was observed for all anti-SARS-CoV-2 specific antibody isotypes and correlated with FcγR binding patterns (**Fig 2C**). These data suggest that IN vaccination induces a higher magnitude and broader antibody subclass response to SARS-CoV-2 than IM vaccination.

**Figure 2.**
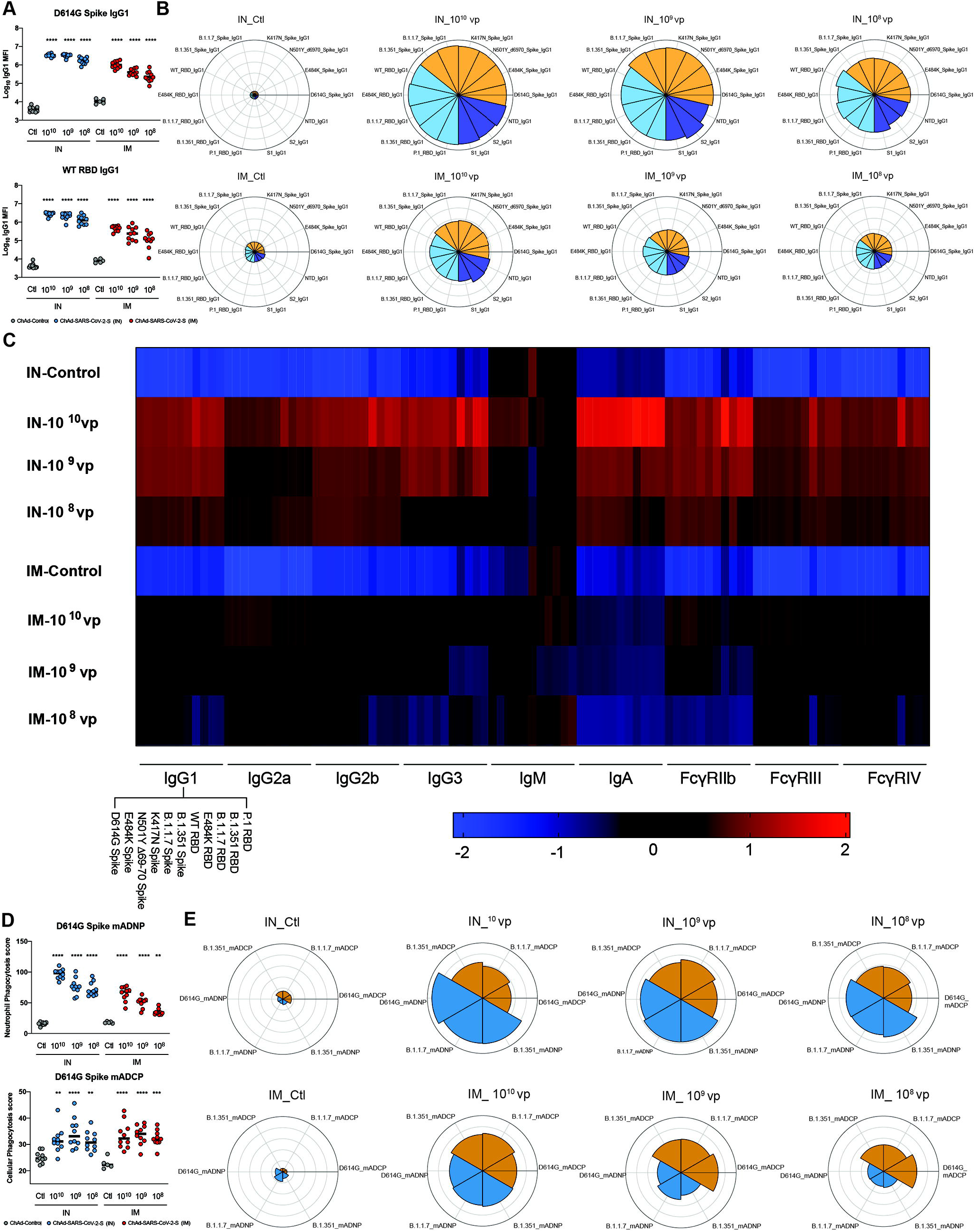
Intranasal inoculation of ChAd-SARS-CoV-2-S induces antibody responses with Fc effector function capacity. **A**. Serum was analyzed by Luminex platform to quantify the amount of anti-SARS-CoV-2 (WA1/2020 D614G) spike and RBD IgG1. Bars represent the mean values. **B**. Serum was analyzed by luminex to quantify the amount of anti-SARS-CoV-2 IgG1 to different SARS-CoV-2 protein variants. Polar plots represent the IgG1 median percentile rank for each SARS-CoV-2 protein and variant. **C**. Heatmap shows the IgG titer and FcγR binding titer of each vaccine regimen to SARS-CoV-2 Spike or RBD proteins. Each square represents the average z-score within a group for the condition. **D**. Serum was incubated with primary mouse neutrophils (mADNP) or J774A.1 cells (mADCP) and SARS-CoV-2 spike-coated beads, and phagocytosis was measured after 1 h. Bars represent the mean and the error bars indicate standard deviations. **E**. Serum was incubated with primary mouse neutrophils (mADNP) or J774A.1 cells (mADCP) and WA1/2020 D614G, B.1.1.7, or B1.351 spike-coated beads, and phagocytosis was measured after 1 h. Polar plots represent the mADNP or mADCP median percentile rank for each SARS-CoV-2 protein and variant. For (**A and D**): one-way ANOVA with a Dunnett’s post-test comparing vaccine to control groups: **, *P* < 0.01; ***, *P* < 0.001; ****, *P* < 0.0001). (**A and D**): Bars indicate median values.

Antibody effector functions, such as opsonization, are mediated in part by Fcγ receptor engagement ^36^. To determine if the observed differences in antibody titers and FcγR binding titers resulted in differences in effector functions, we performed antibody-dependent neutrophil (ADNP) and cellular phagocytosis (ADCP) assays (**Fig 2D-E**). Sera from IN vaccinated mice stimulated substantially more ADNP than those obtained from IM vaccinated mice. However, minimal differences in ADCP were apparent from antibodies derived after IN and IM vaccination (**Fig 2D-E**). These data demonstrate that IN vaccination with ChAd-SARS-CoV-2-S induces a greater and more functional antibody response than after IM vaccination.

### Intranasally administered ChAd-SARS-CoV-2-S induces durable protection against SARS-CoV-2 challenge in BALB/c mice

To assess the efficacy of the ChAd-SARS-CoV-2-S vaccine, immunized BALB/c mice given the dosing regimen described in **Fig 1A** were challenged with SARS-CoV-2. Virus challenge was preceded by intransal introduction of Hu-Ad5-hACE2, which enables ectopic expression of hACE2 and productive infection of SARS-CoV-2 in BALB/c mice by historical SARS-CoV-2 strains ^37, 38^. Animals were immunized once via IN or IM routes with 10^10^ vp of ChAd-Control or 10^8^, 10^9^, or 10^10^ vp of ChAd-SARS-CoV-2-S. At day 95 or 195 post-vaccination, mice were given 10^8^ plaque-forming units (PFU) of Hu-Ad5-hACE2 and anti-Ifnar1 mAb; the latter attenuates innate immunity and enhances pathogenesis in this model ^37^. Five days later, BALB/c mice were challenged with 5 x 10^4^ focus-forming units (FFU) of SARS-CoV-2 (strain WA1/2020) via IN route. At 4 days post-infection (dpi), lungs, spleen, and heart were harvested from mice challenged at 100 days post-immunization, and lungs, nasal turbinates, and nasal washes were collected from a second cohort challenged at 200 days post-immunization. Tissues were assessed for viral burden by quantitative reverse transcription PCR (qRT–PCR) using primers for the subgenomic RNA (N gene). IN immunization with all three doses induced remarkable protection at 100 days post-vaccination as evidenced by a virtual absence of viral RNA in lungs, spleen, and heart compared to animals receiving the ChAd-Control vaccine (**Fig 3A-C**). At 200 days post-immunization, protection conferred by the IN delivered ChAd-SARS-CoV-2-S remained robust in the upper and lower respiratory tracts compared to ChAd-Control immunized mice. Nevertheless, we observed limited infection breakthrough in the lungs and nasal turbinates in animals immunized with the lowest 10^8^ vp dose of ChAd-SARS-CoV-2-S (**Fig 3G and I**). In comparison, protection at 100 days post-IM immunization was less than after IN immunization at the same challenge time point. Although viral RNA was not detected in the heart and spleen (**Fig 3E-F**), at least 1,000 to 30,000-fold (*P* < 0.0001) higher levels were measured in the lungs of mice immunized with ChAd-SARS-CoV-2-S by the IM compared to IN route (**Fig 3A and D**). We also observed a greater impact of dosing by the IM route, as the reduction in viral RNA load in the lung at 10^8^ vp dose no longer was different than in the ChAd-Control vaccinated mice (**Fig 3D**). At 200 days post-IM immunization, we observed less protection against SARS-CoV-2 infection in the lungs, nasal washes, and nasal turbinates than after IN immunization (**Fig 3G-L**).

**Figure 3.**
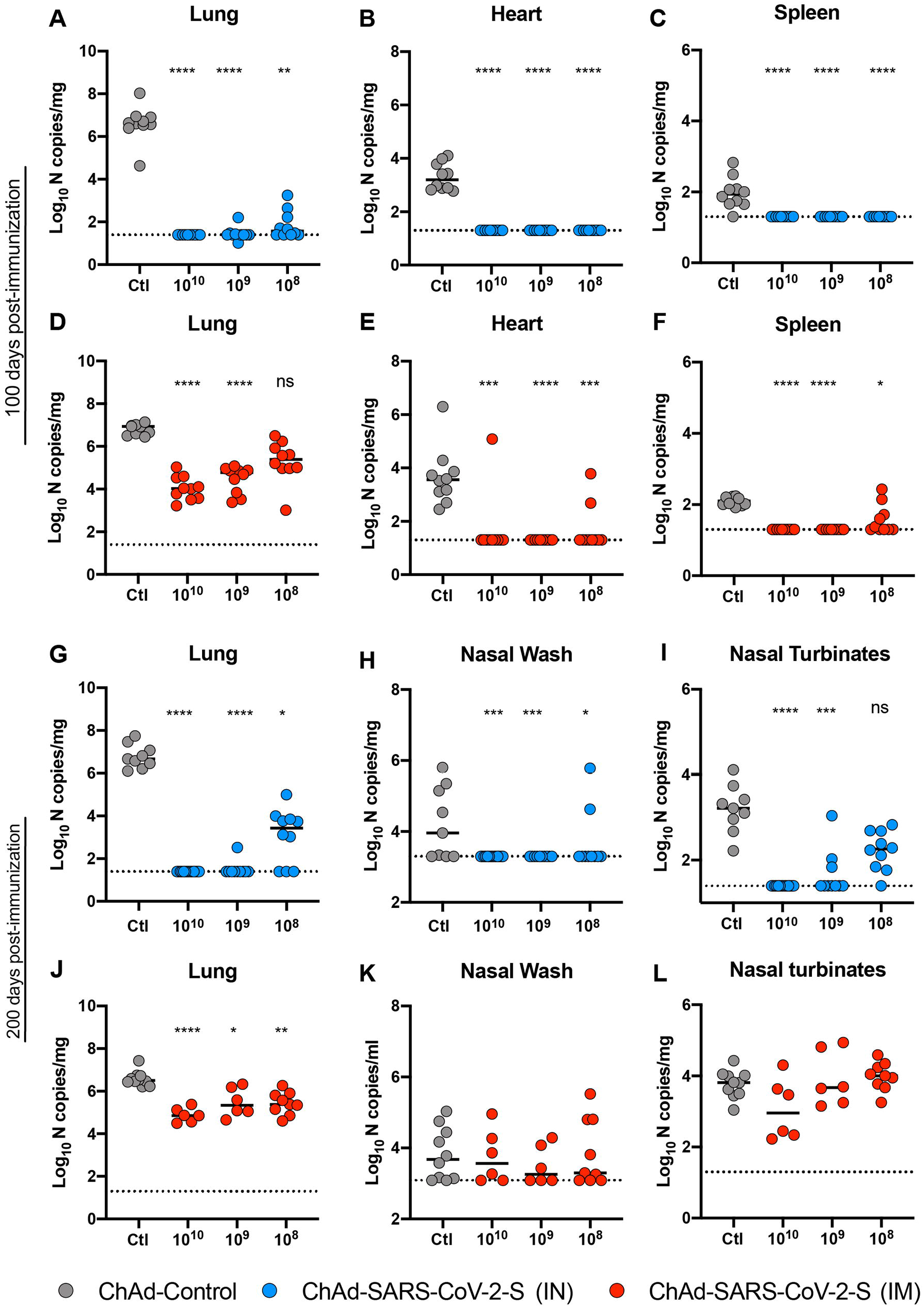
Durability of protective efficacy of ChAd-SARS-CoV-2-S against SARS-CoV-2 infection in BALB/c mice. Five-week old female BALB/c mice were immunized via IN or IM route with 10^10^ vp of ChAd-control or 10^10^, 10^9^ and 10^8^ vp of ChAd-SARS-CoV-2-S. On day 100 or 200 post-immunization, mice were challenged as follows: animals were treated with anti-Ifnar1 mAb and transduced with Hu-AdV5-hACE2 via an IN route one day later. Five days later, mice were inoculated with 5 x 10^4^ FFU of SARS-CoV-2 WA1/2020 via the intranasal route. Tissues were harvested at 4 dpi, and viral RNA levels were measured from mice challenged 100 (**A-F**) or 200 (**G-L**) days post-immunization by RT-qPCR (n = 6-14, Kruskal Wallis with Dunn’s post-test: ns, not significant; **, *P* < 0.01; *, *P* < 0.1; ***, *P* < 0.001 ****, *P* < 0.0001). Bars show median values, and dotted lines indicate the LOD of the assays.

### ChAd-SARS-CoV-2-S induces durable immunity in hACE2 transgenic mice

We next assessed the immunogenicity of intransally-delivered ChAd-SARS-CoV-2-S in K18-hACE2 C57BL/6 mice, which are more vulnerable to SARS-CoV-2 infection than BALB/c mice ^39, 40, 41^. Five-week old K18-hACE2 mice were inoculated via an IN route with 10^9^ vp of ChAd-control or ChAd-SARS-CoV-2-S. Serum samples were collected six weeks later, and humoral immune responses were evaluated. IN immunization of ChAd-SARS-CoV-2-S but not ChAd-control induced high levels of S- and RBD-specific IgG and IgA (**Fig 4A-B**). Neutralizing antibody titers against WA1/2020 and two other SARS-CoV-2 strains with spike proteins from B.1.351 and B.1.1.28 variants were measured by FRNT assay (**Fig 4C-D and S2**). High levels of neutralizing antibody against WA1/2020 were induced after a single IN dose of ChAd-SARS-CoV-2-S (EC50 of 9,591). As seen with vaccine-induced human sera ^24, 25, 26, 27, 28, 29^, we observed decreases in neutralizing titers against Wash-B.1.351 (∼5-fold, *P* < 0.0001; **Fig 4C**) and Wash-B.1.1.28 (∼3-fold, *P* < 0.0001; **Fig. 4D**) SARS-CoV-2 strains compared to WA1/2020. To assess the durability of humoral responses, a separate cohort of K18-hACE2 mice were immunized via the IN route, and serum samples were collected at nine months. ChAd-SARS-CoV-2-S induced high levels of S-and RBD-specific IgG and IgA and neutralizing antibody against WA1/2020 (EC50 of 12,550) at this time point (**Fig 4E-H and S2**). When tested against the Wash-B.1.351 and Wash-B.1.1.28 viruses, we also observed a decrease in neutralizing titer (∼6 to 8-fold, *P* < 0.05; **Fig 4G-H**) compared to WA1/2020, although they still remained high (EC50 of 1,627 and 1,918, respectively).

**Figure 4.**
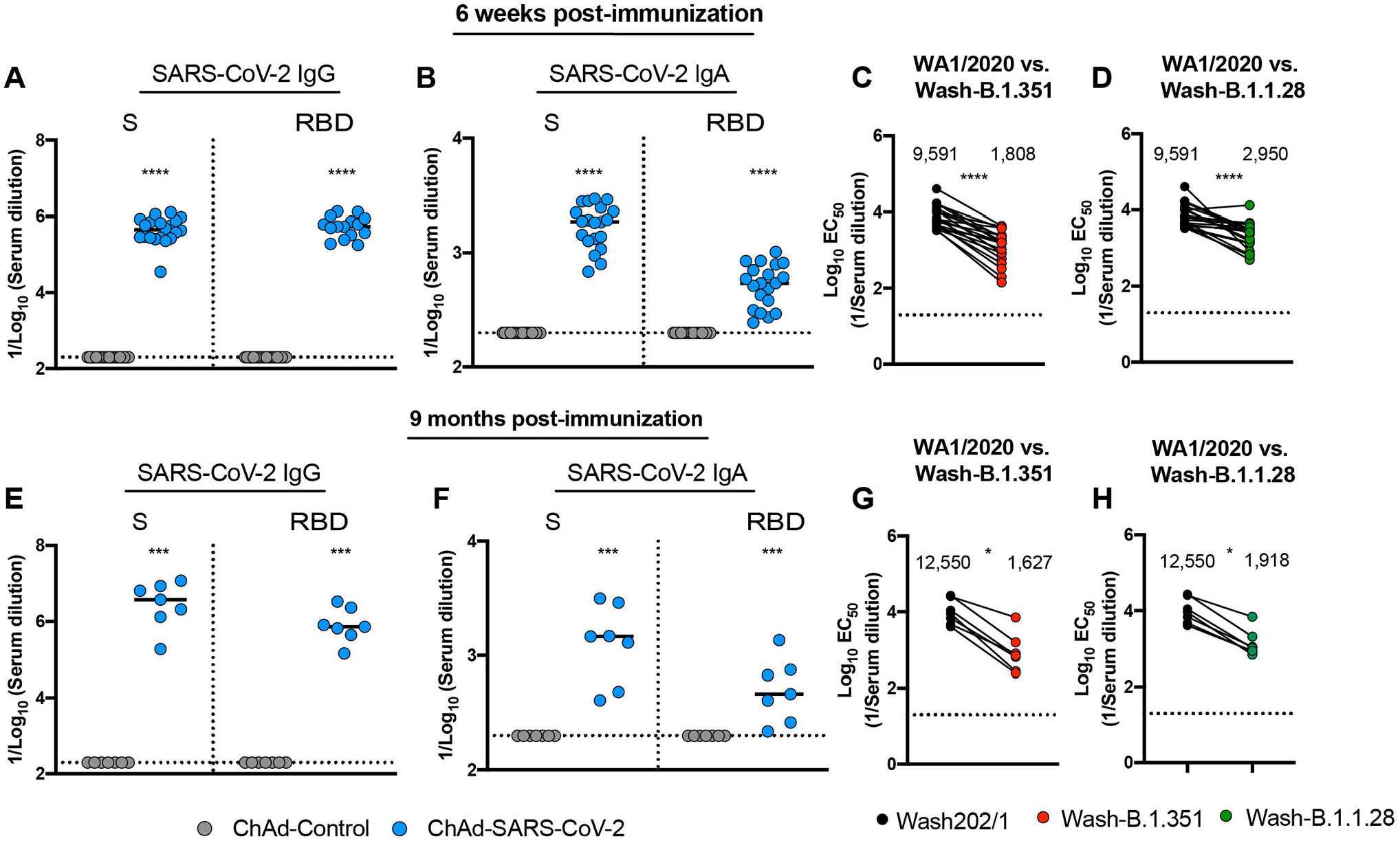
Immunogenicity of intranasally administration ChAd-SARS-CoV-2-S in K18-hACE2 mice. Five-week-old K18-hACE2 female mice were immunized with 10^9^ vp ChAd-control or ChAd-SARS-CoV-2-S via an IN route. Antibody responses in sera of mice at 6 weeks (**A-D**, n = 20) or 9 months (**E-H**, n = 7) after immunization were evaluated. An ELISA measured SARS-CoV-2 S-and RBD-specific IgG (**A, E**) and IgA levels (**B, F**), and an FRNT determined neutralization activity (**C, D, G, H**). **C-D, G-H**, Paired analysis of serum neutralizing activity from immunized mice collected at 6 weeks (**C, D**) or 9 months (**G**) against WA1/2020 and Wash-B.1.351 (**C, G**), or Wash-B.1.1.28 (**D, H**). **A-B, E-F**: Mann-Whitney test: ***, *P* < 0.001; ****, *P* < 0.0001. **C-D, G-H**: Two-tailed Wilcoxon matched-pairs signed rank test: *, *P* < 0.05; ****, *P* < 0.0001.

### ChAd-SARS-CoV-2-S confers cross-protection against Wash B.1.351 and Wash-B.1.1.28 challenge in hACE2 transgenic mice

We tested the protective efficacy of ChAd-SARS-CoV-2-S against WA1/2020 and two chimeric viruses (Wash-B.1.351 and Wash-B.1.1.28) with spike genes corresponding to variants of concern (**Fig 5A**). Five-week old K18-hACE2 mice were immunized via an IN route with a single 10^9^ vp dose of ChAd-control or ChAd-SARS-CoV-2-S. Six weeks later, mice were challenged by an IN route with 10^4^ FFU of Wash-B.1.351, Wash B.1.1.28, or WA1/2020. All mice immunized with ChAd-SARS-CoV-2 exhibited no weight loss, whereas most ChAd-Control-vaccinated mice experienced substantial weight loss at 3 to 6 dpi (**Fig 5B, G, and L**). Remarkably, vaccination with ChAd-SARS-CoV-2-S resulted in almost no detectable SARS-CoV-2 RNA in the upper and lower respiratory tract, heart, and brain at 6 dpi (**Fig 5C-F, H-K, and M-O**). As a further test of the durability of the cross-protective response, five-week old K18-hACE2 mice were immunized via an IN route with a single 10^10^ vp dose of of ChAd-control or ChAd-SARS-CoV-2-S. Nine months later, mice were challenged via IN route with 10^4^ FFU of Wash-B.1.351. ChAd-SARS-CoV-2-S-vaccinated mice maintained weight in contrast to ChAd-Control treated mice (**Fig 5P**). Moreover, substantial virological protection was observed, as only very low amounts of Wash-B.1.351 SARS-CoV-2 RNA were detected in the upper and lower respiratory tracts, heart, and brain in some of the mice (**Fig 5Q-T**).

**Figure 5.**
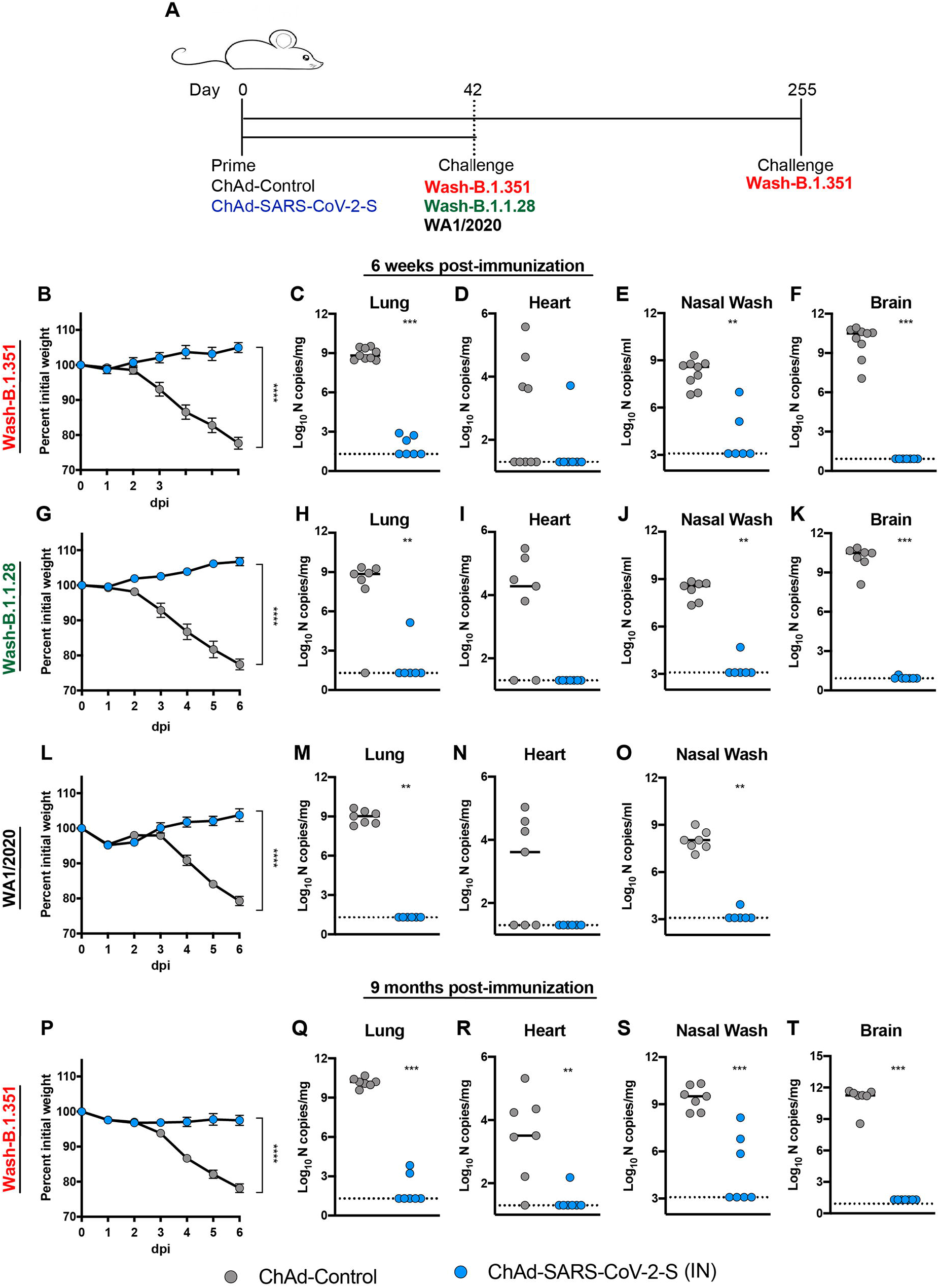
ChAd-SARS-CoV-2-S confers cross-protection aginst variant viruses in K18-hACE2 mice. **A**. Experimental scheme. Five-week-old K18-hACE2 female mice were immunized via an IN route with 10^10^ vp of ChAd-control or ChAd-SARS-CoV-2-S. **B-O**. At 6 weeks post-immunization, mice were challenged with 10^4^ FFU of SARS-CoV-2 of Wash-B.1.351 (**B-F**), Wash-B.1.1.28 (**G-K**), or WA1/2020 (**L-O**). **P-T**. At 9 months post-immunization, mice were challenged with 10^4^ FFU of Wash-B.1.351. **B, G, L, P**. Body weight change over time. Data are the mean ± SEM comparing vaccine to control groups (n = 6-9 for each group; unpaired t test for area under curve, **** *P* < 0.0001). **C-F, H-K, M-O, Q-T**. Viral RNA levels in the lung, heart, brain, and nasal washes were measured at 6 dpi by RT-qPCR (n = 6-9; Mann-Whitney test: ** *P* < 0.01, *** *P* < 0.001). Bars show median values, and dotted lines indicate the LOD of the assays.

## DISCUSSION

The durability of vaccine-induced immune responses is a key for providing sustained protection against SARS-CoV-2 infection and curtailing the current pandemic. Here, we show that a single IN immunization with ChAd-SARS-CoV-2-S induced S-and RBD-specific binding and neutralizing antibodies that continued to rise for several months, suggestive of sustained germinal center reactions. LLPCs in the bone marrow were detected six months after IN vaccination, secreting SARS-CoV-2-specifc IgG and IgA that likely contributed to the durably high antiviral antibody levels in circulation ^34^. In comparison, IM immunization with ChAd-SARS-CoV-2-S induced lower levels of serum neutralizing antibodies, fewer spike-specific IgG secreting LLPCs, and virtually no serum or cellular IgA response. At least in mice, a single IN dose immunization with ChAd-SARS-CoV-2-S produced durable humoral immunity that was observed across a 100-fold dose range. These pre-clinical immunogenicity results compare favorably with studies in humans with mRNA vaccines against SARS-CoV-2, which show humoral immunue responses lasting at least several months ^42, 43^. In comparison, the durability of antibody responses after natural SARS-CoV-2 infection can vary considerably ^44, 45^.

A single immunization of ChAd-SARS-CoV-2-S conferred durable protection against SARS-CoV-2 (WA1/2020 strain) challenge in hACE2-tranduced BALB/c mice or K18-hACE2 transgenic C57BL/6 mice at multiple time points through six months. IN immunization in particular provided virtually complete virological protection against upper and lower respiratory tract infection, with only a limited infection breakthrough seen at the 100-fold lower vaccine dose. The abrogation of infection in the upper respiratory tract suggests that IN vaccination could prevent transmission, although corroborating studies are needed in other rodent (*e*.*g*., hamsters or ferret) models better suited to studying this question ^46^. In comparison, IM immunization reduced the viral RNA levels in the lungs but showed substantially less protection against the homolgous WA1/2020 strain in samples from the upper respiratory tract. While many SARS-CoV-2 vaccine candidates from different platforms have demonstrated immunogenicity and protective efficacy in animals models ^21, 23, 47, 48, 49, 50, 51^, to our knowledge, none have established durability or protection against variant viruses. The long-term protection conferred by IN immunization even at 100-fold lower inoculating doses is promising but remains to be validated in human clinical trials with ChAd-SARS-CoV-2-S. If results in mice were recapitulated, dose sparing strategies could enable production of a large number of vaccine doses that could curtail infection and transmission of SARS-CoV-2.

The emergence of SARS-CoV-2 S variants with mutations of amino acids in the receptor binding motif (*e*.*g*., B.1.351 and B.1.1.28) is of concern because of their resistance to the inhibitory activity of many neutralizing antibodies ^25, 27, 28^. Indeed, human sera from subjects vaccinated with BNT162b2 mRNA or ChAdOx1 nCoV-19 (AZD1222) vaccines showed reduced neutralization against B.1.351 ^28, 52, 53^. Concerningly, IM-administered ChAdOx1 nCoV-19 (AZD1222) showed reduced protective efficacy against mild to moderate B.1.351 infection in humans ^53^. In K18-hACE2 transgenic mice, when we compared the immunogenicity of IN-delivered ChAd-SARS-CoV-2-S against WA1/2020 and chimeric SARS-CoV-2 strains expressing B.1.1.28 or B.1.351 spike proteins, we also observed reduced (3 to 8-fold) neutralization of the variant viruses although the titers remained >1,000. At six weeks post IN immunization of ChAd-SARS-CoV-2-S, K18-hACE2 mice were fully protected mice against weight loss and infection in the upper and lower respiratory tracts and brain by WA1/2020, Wash-B.1.351, and Wash-B.1.1.28. Remarkably, in a separate cohort of K18-hACE2 mice challenged nine months after single IN immunization, animals were fully protected against Wash-B.1.351 challenge. Although correlates of protection are not fully established for SARS-CoV-2 vaccines, the high levels of cross-neutralizing antibodies against the variant viruses combined with robust virus-specific systemic and mucosal CD8^+^ T cell responses described previously ^30^ likely contribute to protection. Beyond this, antibody effector functions also might contribute to prevent SARS-CoV-2 infection and disease ^54, 55, 56^. Indeed, we observed enhanced Fc effector functions against SARS-CoV-2 variant proteins in serum derived from IN-delivered ChAd-SARS-CoV-2-S including robust induction of ADNP and ADCP responses.

### Limitations of study

Although a single intranasal administration of ChAd-SARS-CoV-2-S durably protected against SARS-CoV-2 variant replication in the upper and lower respiratory tract even ∼9 months after immunization, we note several limitations in our study. (1) We performed challenge studies in BALB/c mice transduced with hACE2 or C57BL/6 mice expressing an hACE2 transgene. Durability and protection studies will need to be corroborated in hamsters, non-human primates, and ultimately in humans. (2) Although our studies suggest that the mucosal immunity induced by intranasal vaccination could limit SARS-CoV-2 transmission, the use of mice precluded formal respiratory transmission analysis, which is better studied in hamsters and ferrets ^46^. (3) We observed robust protection *in vivo* against viruses displaying B.1.351 and B.1.1.28 spike proteins likely due to the high serum neutralizing antibody titers. Even though neutralizing antibody levels were lower with the variant strains due to mutations at sites in the receptor binding motif, the high starting level against the historical SARS-CoV-2 likely provided a sufficient cushion to overcome this loss in potency. Studies in other animals or with even lower doses of vaccine where neutralizing titers might be lower are needed to determine if the protective phenotype against variants of concern is maintained. (4) Finally, we did not establish the correlate of protection in these studies, as passive antibody transfer or T cell depletions were not performed. Such studies could be performed in follow-up experiments.

In summary, our study shows that IN immunization with ChAd-SARS-CoV-2-S induces robust and durable binding IgG and IgA antibody, neutralizing antibody, Fc effector functions, and LLPC responses against SARS-CoV-2. In mice, a single IN immunization with ChAd-SARS-CoV-2-S confers cross-protection against SARS-CoV-2 strains displaying spike proteins corresponding to B.1.351 and B.1.1.28 variants, even nine months after vaccination. Given the efficacy of preclinical evaluation in multiple animal models ^30, 31, 32^ and the durable protective immunity against variants of concern, IN delivery of ChAd-SARS-CoV-2-S may be a promising platform for preventing SARS-CoV-2 infection, curtailing transmission, and thus, warrants further clinical evaluation in humans.

## ACKNOWLEDGEMENTS

This study was supported by NIH contracts and grants (R01 AI157155, R01 EB026468-02S1, 75N93019C00062, HHSN272201400018C).

## AUTHOR CONTRIBUTIONS

A.O.H., I.P.D., and E.K. generated the vaccines. A.O.H. performed ELISA assays, and analyzed the data. A.O.H. performed neutralization assays. S.R. performed ELISPOT assays. M.J.G., D.Y., and G.A. designed and/or performed the binding and Fc effector function experiments. L.J.A., D.H.F. designed and produced the recombinant S and RBD proteins and antibodies. A.O.H., S.S., and B.Y. performed and evaluated the virological assays. P-Y.S. provided key reagents. M.S.D., D.H.F., G.A., and D.T.C. designed experiments and secured funding. A.O.H., M.J.G. and M.S.D. wrote the initial draft, with the other authors providing editorial comments.

## SUPPLEMENTAL FIGURE LEGENDS

**Figure S1. ChAd-SARS-CoV-2-S vaccine induces neutralizing antibodies as measured by FRNT. Related to Figure 1**. Five-week old female BALB/c mice were immunized via IN or IM route with a single 10^10^, 10^9^, or 10^8^ dose of ChAd-SARS-CoV-2-S. Serum samples from ChAd-SARS-CoV-2-S vaccinated mice were collected at days 100 (**A**) or 200 (**B**) after immunization and assayed for neutralizing activity by FRNT. Serum neutralization curves corresponding to individual mice are shown for the indicated vaccines (n = 6-14 per group). Each point represents the mean of two technical replicates.

**Figure S2. ChAd-SARS-CoV-2-S vaccine induces neutralizing antibodies as measured by FRNT. Related to Figure 4**. Five-week-old K18-hACE2 female mice were immunized with 10^9^ vp ChAd-control or ChAd-SARS-CoV-2-S via an IN route. Serum samples were collected at six weeks (**A**) or nine months (**B**) after immunization and assayed for neutralizing activity against WA1/2020, Wash-B.1.351, or Wash-B.1.1.28 by FRNT. Serum neutralization curves corresponding to individual mice are shown for the indicated vaccines (n = 7-20 per group). Each point represents the mean of two technical replicates.

## METHODS

### Viruses and cells

Vero E6 (CRL-1586, American Type Culture Collection (ATCC), Vero-TMPRSS2 ^57^, Vero (CCL-81, ATCC) and HEK293 (CRL-1573, ATCC) cells were cultured at 37°C in Dulbecco’s Modified Eagle medium (DMEM) supplemented with 10% fetal bovine serum (FBS), 10□mM HEPES pH 7.3, 1□mM sodium pyruvate, 1X non-essential amino acids, and 100□U/ml of penicillin–streptomycin. Vero-TMPRSS2 cells also were supplemented with 5 μg/mL of blasticidin.

SARS-CoV-2 strain 2019n-CoV/USA_WA1/2020 (WA1/2020) was obtained from the Centers for Disease Control and Prevention. The virus was passaged once in Vero CCL-81 cells and titrated by focus-forming assay (FFA) on Vero E6 cells. The Wash-B.1.351 and Wash-B.1.1.28 chimeric viruses with variant spike genes were described previously ^28, 58^. All viruses were passaged once in Vero-TMPRSS2 cells and subjected to next-generation sequencing to confirm the introduction and stability of substitutions. All virus experiments were performed in an approved Biosafety level 3 (BSL-3) facility.

### Mouse experiments

Animal studies were carried out in accordance with the recommendations in the Guide for the Care and Use of Laboratory Animals of the National Institutes of Health. The protocols were approved by the Institutional Animal Care and Use Committee at the Washington University School of Medicine (Assurance number A3381-01). Virus inoculations were performed under anesthesia that was induced and maintained with ketamine hydrochloride and xylazine, and all efforts were made to minimize animal suffering.

Female BALB/c (catalog 000651) and K18-hACE2 C57BL/6 (catalog 034860) mice were purchased from The Jackson Laboratory. Four to five-week-old animals were immunized with 10^10^ vp of ChAdV-control or 10^8^, 10^9^, or 10^10^ vp of ChAd-SARS-CoV-2-S in 50 µl PBS via IM (hind leg) or IN injection. Vaccinated BALB/c mice (10 to 11-week-old) were given a single intraperitoneal injection of 2 mg of anti-Ifnar1 mAb (MAR1-5A3 ^59^ (Leinco) one day before IN administration of 10^8^ PFU of Hu-Ad5-hACE2 ^37^. Five days after Hu-Ad5--hACE2 transduction, mice were inoculated with 4 x 10^5^ FFU of WA1/2020 SARS-CoV-2 by the IN route. K18-hACE2 mice were challenged on indicated days after immunization with 10^4^ FFU of SARS-CoV-2 (WA1/2020, Wash-B.1.351, or Wash-B.1.1.28) via IN route. Animals were euthanized at 6 dpi, and tissues were harvested for virological analysis.

### Chimpanzee and human adenovirus vectors

The ChAd-SARS-CoV-2 and ChAd-Control vaccine vectors were derived from simian Ad36 backbones ^60^, and the constructing and validation has been described in detail previously ^30^. The rescued replication-incompetent ChAd-SARS-CoV-2-S and ChAd-Control vectors were scaled up in HEK293 cells and purified by CsCl density-gradient ultracentrifugation. Viral particle concentration in each vector preparation was determined by spectrophotometry at 260 nm as described ^61^. The Hu-AdV5-hACE2 vector also was described previously ^30^ and produced in HEK293 cells. The viral titer was determined by plaque assay in HEK293 cells.

### SARS-CoV-2 neutralization assays

Heat-inactivated serum samples were diluted serially and incubated with 10^2^ FFU of different SARS-CoV-2 strains for 1 h at 37°C. The virus-serum mixtures were added to Vero cell monolayers in 96-well plates and incubated for 1 h at 37°C. Subsequently, cells were overlaid with 1% (w/v) methylcellulose in MEM supplemented with 2% FBS. Plates were incubated for 30 h before fixation using 4% PFA in PBS for 1 h at room temperature. Cells were washed then sequentially incubated with an oligoclonal pool of SARS2-2, SARS2-11, SARS2-16, SARS2-31, SARS2-38, SARS2-57, and SARS2-71 ^62^ anti-S antibodies and HRP-conjugated goat anti-mouse IgG (Sigma, 12-349) in PBS supplemented with 0.1% saponin and 0.1% bovine serum albumin. TrueBlue peroxidase substrate (KPL) was used to develop the plates before counting the foci on a BioSpot analyzer (Cellular Technology Limited).

### Protein expression and purification

The cloning and production of purified S and RBD proteins corresponding to the WA1/2020 SARS-CoV-2 strain have been described previously ^30, 63^. Briefly, prefusion-stabilized S ^64^ and RBD were cloned into a pCAGGS mammalian expression vector with a hexahistidine tag and transiently transfected into Expi293F cells. Proteins were purified by cobalt-charged resin chromatography (G-Biosciences).

### ELISA

Purified antigens (S or RBD) were coated onto 96-well Maxisorp clear plates at 2 µg/mL in 50 mM Na_2_CO_3_ pH 9.6 (70 µL) overnight at 4°C. Coating buffers were aspirated, and wells were blocked with 200 µL of 1X PBS + 0.05% Tween-20 + 1% BSA + 0.02% NaN_3_ (Blocking buffer, PBSTBA) overnight at 4°C. Heat-inactivated serum samples were diluted in PBSTBA in a separate 96-well polypropylene plate. The plates then were washed thrice with 1X PBS + 0.05% Tween-20 (PBST), followed by addition of 50 µL of respective serum dilutions. Sera were incubated in the blocked ELISA plates for at least 1 h at room temperature. The ELISA plates were again washed thrice in PBST, followed by addition of 50 µL of 1:1,000 anti-mouse IgG-HRP (Southern Biotech Cat. #1030-05) in PBST or 1:1000 of anti-mouse IgA-HRP in PBSTBA (SouthernBiotech). Plates were incubated at room temperature for 1 h, washed thrice in PBST, and then 100 µL of 1-Step Ultra TMB-ELISA was added (ThermoFisher Cat. #34028). Following a 10 to 12-min incubation, reactions were stopped with 50 µL of 2 M sulfuric acid. Optical density (450 nm) measurements were determined using a microplate reader (Bio-Rad).

### ELISPOT assay

To quantitate S-specific plasma cells in the bone marrow, femurs and tibias were crushed using a mortar and pestle in RPMI 1640, filtered through a 100 μm strainer and subjected to ACK lysis. CD138^+^ cells were enriched by positive selection and magnetic beads according to the manufacturer’s instructions (EasySep Mouse CD138 Positive Selection, STEMCELL). The enriched CD138^+^ cells were incubated overnight in RPMI 1640 supplemented with 10% FBS in MultiScreen-HA Filter Plates (Millipore) pre-coated with SARS-CoV-2 S protein. Foci were developed using TruBlue substrate (KPL) following sequential incubation with anti-mouse IgG-biotin or anti-mouse IgA-biotin and streptavidin-HRP. Plates were imaged using a BioSpot instrument, and foci enumerated manually.

### Measurement of viral burden

SARS-CoV-2 infected mice were euthanized using a ketamine and xylazine cocktail, and organs were collected. Tissues were weighed and homogenized with beads using a MagNA Lyser (Roche) in 1 ml of Dulbecco’s Modified Eagle’s Medium (DMEM) containing 2% fetal bovine serum (FBS). RNA was extracted from clarified tissue homogenates using MagMax mirVana Total RNA isolation kit (Thermo Scientific) and the KingFisher Flex extraction system (Thermo Scientific). SARS-CoV-2 RNA levels were measured by one-step quantitative reverse transcriptase PCR (qRT-PCR) TaqMan assay as described previously ^37^. SARS-CoV-2 nucleocapsid (N) specific primers and probe sets were used: (N: F Primer: ATGCTGCAATCGTGCTACAA; R primer: GACTGCCGCCTCTGCTC; probe: /56-FAM/TCAAGGAAC/ZEN/AACATTGCCAA/3IABkFQ) (Integrated DNA Technologies). Viral RNA was expressed as (N) gene copy numbers per milligram on a log_10_ scale.

### Luminex analysis

Luminex analhysis was conducted as described previously ^65^. Briefly, proteins (Spike: D614G, E484K, N501Δ69-70, K417N, B.1.1.7, B.1.351; Receptor Binding Domain (RBD) (ImmuneTech): WT, E484K, B.1.1.7, B.1.351, B.1.128) were carboxy-coupled to magnetic Luminex microplex carboxylated beads (Luminex Corporation) using NHS-ester linkages with Sulfo-NHS and EDC (Thermo Fisher) and then incubated with serum (IgG1, FcγRIIb, FcγRIII 1:3000; IgG2a, G2b, G3, A, FcγRIV 1:1000, IgM 1:500) for 2 h at 37°C. Isotype analysis was perfomed by incubating the immune complexes with secondary goat anti-mouse-PE antibody (IgG1 1070-09, IgG2a 1080-09S, IgG2b 1090-09S, IgG3 1100-09, IgM 1020-09, IgA 1040-09 Southern Biotech) for each isotype. FcγR binding was quantified by incubating immune complexes with biotinylated FcγRs (FcγRIIB, FcγRIII, and FcγRIV, courtesy of Duke Protein Production Facility) conjugated to Steptavidin-PE (Prozyme). Flow cytometry was performed with an IQue (Intellicyt) and analysis was performed on IntelliCyt ForeCyt (v8.1).

### Antibody-dependent neutrophil or cellular phagocytosis

Antibody-dependent neutrophil phagocytosis (ADNP) and cellular phagocytosis (ADCP) assays were conducted as described previously ^66, 67, 68^. Briefly, spike protein was carboxy coupled to blue, yellow-green, or red FluoSphere™ Carboxylate-modified microsphere, 0.2 μm (ThermoFisher) using NHS-ester linkages with Sulfo-NHS and EDC (Thermo Fisher). Spike-coated beads were incubated with diluted serum (1:150 ADNP, 1:100 ADCP) for 2 hours at 37°C. For the ADNP assay, bone marrow cells were collected from BALB/c mice, and red blood cells were subjected to ACK lysis. The remaining cells were washed with PBS, and aliquoted into 96-well plates (5 x 10^4^ cells per well). The bead-antibody complexes were added to cells and incubated for 1 h at 37°C. After washing, cells were stained with the following antibodies: CD11b APC (BioLegend 101212), CD11c A700 (BioLegend 117320), Ly6G Pacific Blue (127628), Ly6C BV605 (BioLegend 128036), Fcblock (BD Bioscience 553142) and CD3 PE/Cy7 (BioLegend 100320). Cells were fixed with 4% PFA, processed on an BD LSRFortessa (BD Biosciences). Neutrophils were defined as CD3^-^, CD11b^+^, Ly6G^+^. The neutrophil phagocytosis score was calculated as (% FITC+) x (geometic mean fluorescent intensity of FITC)/10000. For the ADCP assay, J774A.1 (ATCC TIB-67) murine monocytic cells were incubated with the Spike-coated bead–antibody complexes for 1□h at 37°C. Cells were washed in 5□mM EDTA PBS, fixed with 4% PFA, and analyzed on an BD LSRFortessa (BD Biosciences). The cellular phagocytosis score was calculated as (% FITC+) x (geometic mean fluorescent intensity of FITC)/10000.

### Statistical analysis

Statistical significance was assigned when *P* values were < 0.05 using Prism Version 8 (GraphPad) or Jupyter Notebook 6.1.4. Tests, number of animals (n), median values, and statistical comparison groups are indicated in the Figure legends. Analysis of anti-S, anti-RBD, neutralization titers, and ELISPOT values after vaccination was performed using a one-way ANOVA with a Dunnett’s post-test or a Mann-Whitney test. Differences in viral titers after SARS-CoV-2 infection of immunized mice were determined using a Kruskal Wallis ANOVA with Dunn’s post-test or a Mann-Whitney test. Differences in neutralization titers for different variants were compared using two-tailed Wilcoxon matched-pairs signed rank test. Weight changes were analyzed using area under the curve analysis and a Student’s t test.

## Data and code availability

All data supporting the findings of this study are available within the paper and are available from the corresponding author upon request.

## REFERENCES

1. Mao, R. et al. Manifestations and prognosis of gastrointestinal and liver involvement in patients with COVID-19: a systematic review and meta-analysis. The lancet. Gastroenterology & hepatology (2020).

2. Wichmann, D. et al. Autopsy Findings and Venous Thromboembolism in Patients With COVID-19: A Prospective Cohort Study. Ann Intern Med 173, 268–277 (2020).

3. Cheung, E.W. et al. Multisystem Inflammatory Syndrome Related to COVID-19 in Previously Healthy Children and Adolescents in New York City. JAMA 324, 294–296 (2020).

4. Zhou, F. et al. Clinical course and risk factors for mortality of adult inpatients with COVID-19 in Wuhan, China: a retrospective cohort study. Lancet 395, 1054–1062 (2020).

5. Letko, M., Marzi, A. & Munster, V. Functional assessment of cell entry and receptor usage for SARS-CoV-2 and other lineage B betacoronaviruses. Nature microbiology 5, 562–569 (2020).

6. Hoffmann, M. et al. SARS-CoV-2 Cell Entry Depends on ACE2 and TMPRSS2 and Is Blocked by a Clinically Proven Protease Inhibitor. Cell (2020).

7. Wrapp, D. et al. Cryo-EM structure of the 2019-nCoV spike in the prefusion conformation. Science 367, 1260–1263 (2020).

8. Pinto, D. et al. Cross-neutralization of SARS-CoV-2 by a human monoclonal SARS-CoV antibody. Nature 583, 290–295 (2020).

9. Cao, Y. et al. Potent neutralizing antibodies against SARS-CoV-2 identified by high-throughput single-cell sequencing of convalescent patients’ B cells. Cell 182, 73–84 (2020).

10. Zost, S.J. et al. Rapid isolation and profiling of a diverse panel of human monoclonal antibodies targeting the SARS-CoV-2 spike protein. Nat Med 26, 1422–1427 (2020).

11. Barnes, C.O. et al. SARS-CoV-2 neutralizing antibody structures inform therapeutic strategies. Nature 588, 682–687 (2020).

12. Tortorici, M.A. et al. Ultrapotent human antibodies protect against SARS-CoV-2 challenge via multiple mechanisms. Science 370, 950–957 (2020).

13. Cao, L. et al. De novo design of picomolar SARS-CoV-2 miniprotein inhibitors. Science 370, 426–431 (2020).

14. Burton, D.R. & Walker, L.M. Rational Vaccine Design in the Time of COVID-19. Cell Host Microbe 27, 695–698 (2020).

15. Graham, B.S. Rapid COVID-19 vaccine development. Science 368, 945–946 (2020).

16. Polack, F.P. et al. Safety and Efficacy of the BNT162b2 mRNA Covid-19 Vaccine. N Engl J Med 383, 2603–2615 (2020).

17. Baden, L.R. et al. Efficacy and Safety of the mRNA-1273 SARS-CoV-2 Vaccine. N Engl J Med 385, 403–416 (2021).

18. Sadoff, J. et al. Interim Results of a Phase 1-2a Trial of Ad26.COV2.S Covid-19 Vaccine. N Engl J Med (2021).

19. Barrett, J.R. et al. Phase 1/2 trial of SARS-CoV-2 vaccine ChAdOx1 nCoV-19 with a booster dose induces multifunctional antibody responses. Nat Med 27, 279–288 (2021).

20. Mercado, N.B. et al. Single-shot Ad26 vaccine protects against SARS-CoV-2 in rhesus macaques. Nature 586, 583–588 (2020).

21. van Doremalen, N. et al. ChAdOx1 nCoV-19 vaccine prevents SARS-CoV-2 pneumonia in rhesus macaques. Nature 586, 578–582 (2020).

22. Wang, H. et al. Development of an Inactivated Vaccine Candidate, BBIBP-CorV, with Potent Protection against SARS-CoV-2. Cell 182, 713-721.e719 (2020).

23. Yu, J. et al. DNA vaccine protection against SARS-CoV-2 in rhesus macaques. Science 369, 806–811 (2020).

24. Wibmer, C.K. et al. SARS-CoV-2 501Y.V2 escapes neutralization by South African COVID-19 donor plasma. bioRxiv (2021).

25. Wang, Z. et al. mRNA vaccine-elicited antibodies to SARS-CoV-2 and circulating variants. Nature (2021).

26. Tada, T. et al. Neutralization of viruses with European, South African, and United States SARS-CoV-2 variant spike proteins by convalescent sera and BNT162b2 mRNA vaccine-elicited antibodies. bioRxiv (2021).

27. Wang, P. et al. Antibody Resistance of SARS-CoV-2 Variants B.1.351 and B.1.1.7. Nature (2021).

28. Chen, R.E. et al. Resistance of SARS-CoV-2 variants to neutralization by monoclonal and serum-derived polyclonal antibodies. Nat Med (2021).

29. McCallum, M. et al. SARS-CoV-2 immune evasion by variant B.1.427/B.1.429. bioRxiv (2021).

30. Hassan, A.O. et al. A Single-Dose Intranasal ChAd Vaccine Protects Upper and Lower Respiratory Tracts against SARS-CoV-2. Cell 183, 169-184.e113 (2020).

31. Bricker, T.L. et al. A single intranasal or intramuscular immunization with chimpanzee adenovirus vectored SARS-CoV-2 vaccine protects against pneumonia in hamsters. bioRxiv (2020).

32. Hassan, A.O. et al. A single intranasal dose of chimpanzee adenovirus-vectored vaccine protects against SARS-CoV-2 infection in rhesus macaques. Cell Reports Medicine (2021).

33. Case, J.B. et al. Neutralizing antibody and soluble ACE2 inhibition of a replication-competent VSV-SARS-CoV-2 and a clinical isolate of SARS-CoV-2. Cell Host and Microbe 28, 475–485 (2020).

34. Amanna, I.J. & Slifka, M.K. Mechanisms that determine plasma cell lifespan and the duration of humoral immunity. Immunol Rev 236, 125–138 (2010).

35. Purtha, W.E., Tedder, T.F., Johnson, S., Bhattacharya, D. & Diamond, M.S. Memory B cells but not long-lived plasma cells possess antigen specificities for viral escape mutants J Exp Med 208, 2599–2606 (2011).

36. Bruhns, P. & Jönsson, F. Mouse and human FcR effector functions. Immunol Rev 268, 25–51 (2015).

37. Hassan, A.O. et al. A SARS-CoV-2 Infection Model in Mice Demonstrates Protection by Neutralizing Antibodies. Cell (2020).

38. Sun, S.H. et al. A Mouse Model of SARS-CoV-2 Infection and Pathogenesis. Cell Host Microbe (2020).

39. Winkler, E.S. et al. SARS-CoV-2 infection of human ACE2-transgenic mice causes severe lung inflammation and impaired function. Nat Immunol 21, 1327–1335 (2020).

40. Yinda, C.K. et al. K18-hACE2 mice develop respiratory disease resembling severe COVID-19. PLoS Pathog 17, e1009195 (2021).

41. Golden, J.W. et al. Human angiotensin-converting enzyme 2 transgenic mice infected with SARS-CoV-2 develop severe and fatal respiratory disease. JCI insight 5 (2020).

42. Widge, A.T. et al. Durability of Responses after SARS-CoV-2 mRNA-1273 Vaccination. N Engl J Med 384, 80–82 (2021).

43. Doria-Rose, N. et al. Antibody Persistence through 6 Months after the Second Dose of mRNA-1273 Vaccine for Covid-19. N Engl J Med (2021).

44. Gudbjartsson, D.F. et al. Humoral Immune Response to SARS-CoV-2 in Iceland. N Engl J Med 383, 1724–1734 (2020).

45. Dan, J.M. et al. Immunological memory to SARS-CoV-2 assessed for up to 8 months after infection. Science (2021).

46. Muñoz-Fontela, C. et al. Animal models for COVID-19. Nature 586, 509–515 (2020).

47. Vogel, A.B. et al. BNT162b vaccines protect rhesus macaques from SARS-CoV-2. Nature 592, 283–289 (2021).

48. García-Arriaza, J. et al. COVID-19 vaccine candidates based on modified vaccinia virus Ankara expressing the SARS-CoV-2 spike induce robust T- and B-cell immune responses and full efficacy in mice. J Virol (2021).

49. Yao, Y.F. et al. Protective Efficacy of Inactivated Vaccine against SARS-CoV-2 Infection in Mice and Non-Human Primates. Virologica Sinica, 1–11 (2021).

50. Hennrich, A.A. et al. Safe and effective two-in-one replicon-and-VLP minispike vaccine for COVID-19: Protection of mice after a single immunization. PLoS Pathog 17, e1009064 (2021).

51. Tostanoski, L.H. et al. Ad26 vaccine protects against SARS-CoV-2 severe clinical disease in hamsters. Nat Med (2020).

52. Zhou, D. et al. Evidence of escape of SARS-CoV-2 variant B.1.351 from natural and vaccine-induced sera. Cell (2021).

53. Madhi, S.A. et al. Efficacy of the ChAdOx1 nCoV-19 Covid-19 Vaccine against the B.1.351 Variant. N Engl J Med (2021).

54. Winkler, E.S. et al. Human neutralizing antibodies against SARS-CoV-2 require intact Fc effector functions for optimal therapeutic protection. Cell 184, 1804-1820.e1816 (2021).

55. Schäfer, A. et al. Antibody potency, effector function, and combinations in protection and therapy for SARS-CoV-2 infection in vivo. J Exp Med 218 (2021).

56. Bartsch, Y.C. et al. Discrete SARS-CoV-2 antibody titers track with functional humoral stability. Nat Commun 12, 1018 (2021).

57. Zang, R. et al. TMPRSS2 and TMPRSS4 promote SARS-CoV-2 infection of human small intestinal enterocytes. Sci Immunol 5 (2020).

58. Xie, X. et al. An Infectious cDNA Clone of SARS-CoV-2. Cell Host Microbe 27, 841-848.e843 (2020).

59. Sheehan, K.C. et al. Blocking monoclonal antibodies specific for mouse IFN-alpha/beta receptor subunit 1 (IFNAR-1) from mice immunized by in vivo hydrodynamic transfection. J Interferon Cytokine Res 26, 804–819 (2006).

60. Roy, S. et al. Creation of a panel of vectors based on ape adenovirus isolates. J Gene Med 13, 17–25 (2011).

61. Maizel, J.V., Jr., White, D.O. & Scharff, M.D. The polypeptides of adenovirus. I. Evidence for multiple protein components in the virion and a comparison of types 2, 7A, and 12. Virology 36, 115–125 (1968).

62. Liu, Z. et al. Identification of SARS-CoV-2 spike mutations that attenuate monoclonal and serum antibody neutralization. Cell Host Microbe 29, 477-488.e474 (2021).

63. Alsoussi, W.B. et al. A Potently Neutralizing Antibody Protects Mice against SARS-CoV-2 Infection. J Immunol (2020).

64. Hsieh, C.L. et al. Structure-based design of prefusion-stabilized SARS-CoV-2 spikes. Science 369, 1501–1505 (2020).

65. Brown, E.P. et al. Multiplexed Fc array for evaluation of antigen-specific antibody effector profiles. J Immunol Methods 443, 33–44 (2017).

66. Wessel, A.W. et al. Antibodies targeting epitopes on the cell-surface form of NS1 protect against Zika virus infection during pregnancy. Nat Commun 11, 5278 (2020).

67. Gunn, B.M. et al. A Fc engineering approach to define functional humoral correlates of immunity against Ebola virus. Immunity 54, 815-828.e815 (2021).

68. Butler, A.L., Fallon, J.K. & Alter, G. A Sample-Sparing Multiplexed ADCP Assay. Front Immunol 10, 1851 (2019).

