## Supplementary figures and images for "An intranasal vaccine durably protects against SARS-CoV-2 variants in mice"

### Supplemental Figure S2

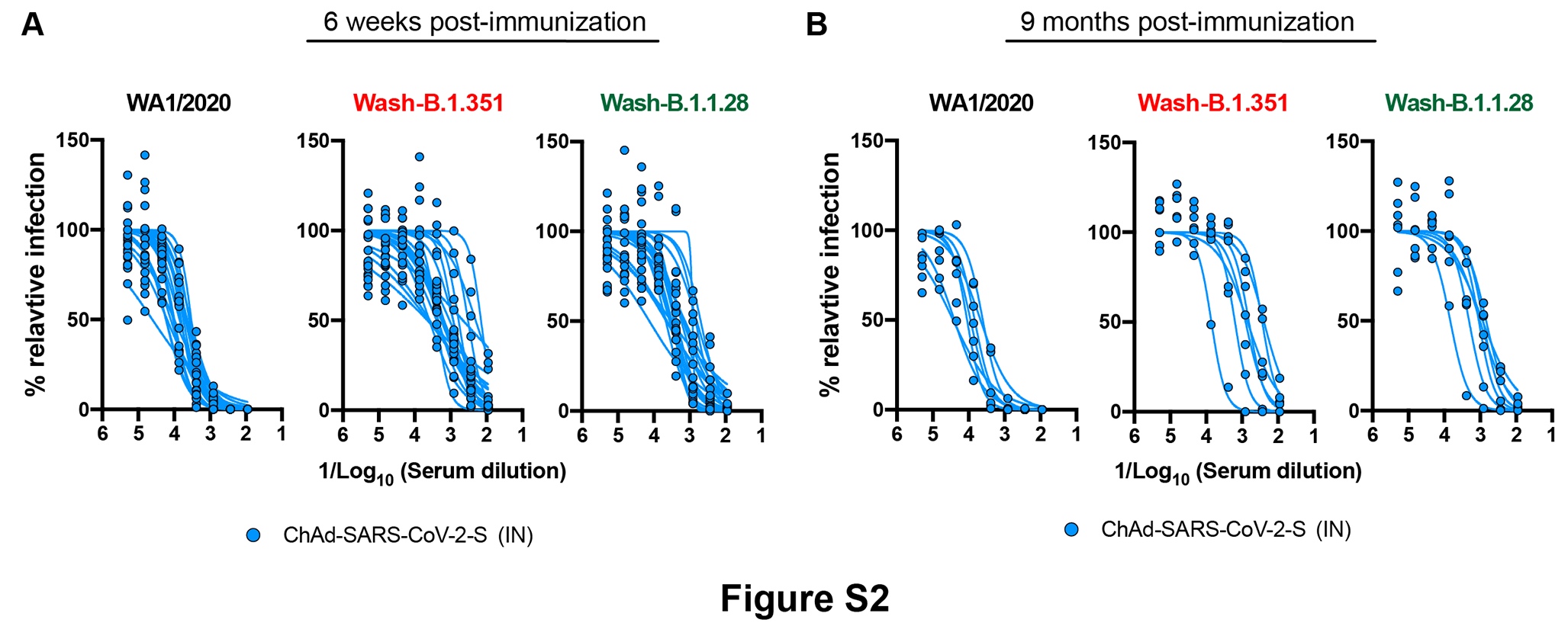

### Supplemental Figures S1

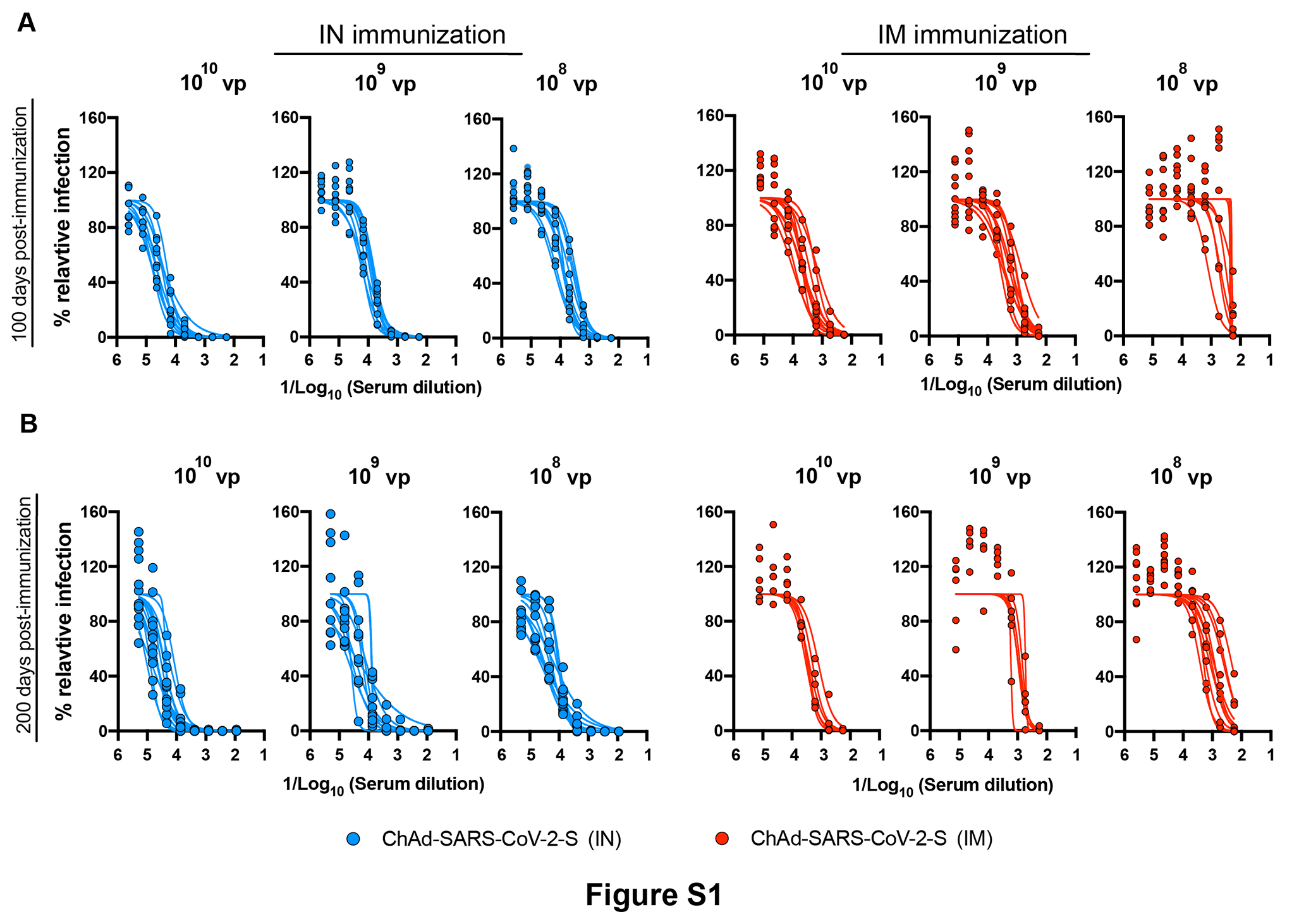
